# Spinal V1 neurons inhibit motor targets locally and sensory targets distally to coordinate locomotion

**DOI:** 10.1101/2021.01.22.427836

**Authors:** Mohini Sengupta, Vamsi Daliparthi, Yann Roussel, Tuan Vu Bui, Martha W. Bagnall

## Abstract

Rostro-caudal coordination of spinal motor output is essential for locomotion. Most spinal interneurons project axons longitudinally to govern locomotor output, yet their connectivity along this axis remains unclear. In this study, we use larval zebrafish to map synaptic outputs of a major inhibitory population, V1 (Eng1+) neurons, which are implicated in dual sensory and motor functions. We find that V1 neurons exhibit long axons extending rostrally and exclusively ipsilaterally for an average of 6 spinal segments; however, they do not connect uniformly with their post-synaptic targets along the entire length of their axon. Locally, V1 neurons inhibit motor neurons (both fast and slow) and other premotor targets including V2a, V2b and commissural pre-motor neurons. In contrast, V1 neurons make robust inhibitory contacts throughout the rostral extent of their axonal projections onto a dorsal horn sensory population, the Commissural Primary Ascending neurons (CoPAs). In a computational model of the ipsilateral spinal network, we show that this pattern of short range V1 inhibition to motor and premotor neurons is crucial for coordinated rostro-caudal propagation of the locomotor wave. We conclude that spinal network architecture in the longitudinal axis can vary dramatically, with differentially targeted local and distal connections, yielding important consequences for function.

## Introduction

The structure of neuronal connectivity is key to function. In cortex, the structure of inhibitory circuits influences synaptic gain, spike timing, and membrane potential oscillations^1^. In vertebrates, the spinal cord is a major seat for motor control as it contains local circuits necessary and sufficient for producing movement. Like the brain, the spinal cord also contains a range of distinct interneuron classes, the interplay of which results in a rich repertoire of movements^2^. Extensive work has focused on understanding the functions of these interneuron classes evaluating the behavioral effects of genetic ablation^3–6^; however, at the mechanistic level, the circuit architecture underlying these effects remains largely unclear. Defining spinal network organization can help provide valuable insight into interneuron functions^7–11^ especially the spatial structure of connectivity^12^.

The spinal cord is elongated in the longitudinal or rostro-caudal (R-C) axis, which consists of many repeated segments. Coordination along this axis is crucial for locomotion^13,14^, yet organization of neurons in this dimension is poorly understood. Based on studies of transected spinal preparations, coordination in the R-C axis is not set up by independent segmental circuits, but instead relies on continuous, segment spanning networks^15^ including long propriospinal interneurons. Interestingly, blockade of glycinergic neurons disrupts R-C coordination independently of left-right alternation, implying that ipsilateral inhibition is vital for locomotor propagation^15^. Most spinal interneurons, including ipsilateral inhibitory neurons, project axons spanning several segments along the R-C axis^16–20^ yet not much is known about their connectivity in this axis or its functional implications. Here, taking advantage of the transparency and accessibility of the intact spinal cord in larval zebrafish, as well as its significant homology with other vertebrates, we mapped connectivity along the R-C axis in a major ipsilateral inhibitory population: V1 neurons.

V1 interneurons are marked by the expression of Engrailed1 (Eng1) transcription factor across vertebrates^18,21,22^. Genetic ablation of these neurons reduces locomotor speeds in both zebrafish^3^ and mice^6,23^ indicating that speed regulation is a primitive function of these neurons. In limbed vertebrates, V1 serve as Renshaw cells to provide recurrent inhibition onto motor neurons, and as Ia inhibitory neurons to provide reciprocal inhibition enforcing flexor-extensor alternation ^24–26^. Interestingly, connectivity studies have also revealed that V1 neurons inhibit sensory targets, suggesting yet another role for these neurons: sensory gating during locomotion^18,22^. It is unknown whether and how the motor and sensory functions of V1 neurons are organized along the longitudinal axis of the spinal cord.

Using a combination of single cell labelling, optogenetics and electrophysiology in vivo, we mapped synaptic connectivity from V1 neurons to eight motor and sensory spinal populations. Our results reveal that V1 neurons exhibit differential connectivity as they traverse the spinal cord longitudinally. Despite projecting long axons spanning > 5 segments, V1 neurons inhibit motor targets only locally. In contrast, they inhibit sensory targets distally. Using our connectivity map as the basis of a simplified model of the ipsilateral spinal cord, we show that this structure of V1 inhibition is critical for maintaining R-C coordination and locomotor speed.

## Materials and Methods

### Animals

Adult zebrafish (Danio rerio) were maintained at 28.5°C with a 14:10 light:dark cycle in the Washington University Zebrafish Facility following standard care procedures. Larval zebrafish, 4–6 days post fertilization (dpf), were used for experiments and kept in petri dishes in system water or housed with system water flow. Animals older than 5 dpf were fed rotifers or dry powder daily. All procedures described in this work adhere to NIH guidelines and received approval by the Washington University Institutional Animal Care and Use Committee.

### Transgenic Fish Lines

The transgenic line *Tg(eng1b-hs:Gal4)nns40Tg* (ZDB-ALT-151202-14) was generated from CRISPR-mediated transgenesis^27^ and was a kind gift from Dr. Shin-ichi Higashijima. This line was crossed with a stable *Tg(UAS:CatCh)stl602* line (ZDB-ALT-201209-12) generated by our lab^28^ to create double transgenic animals that we refer to throughout this paper as *Tg(eng1b:Gal4,UAS:CatCh*). For targeting V2a and V2b neurons, the *Tg(vsx2:loxP-DsRed-loxP-GFP*)*nns3Tg* (ZDB-ALT-061204-4)^29^, and *Tg(gata3:lox-Dsred-lox:GFP)nns53Tg* (ZDB-ALT-190724-4)^28^ lines, respectively, were crossed to *Tg(eng1b:Gal4,UAS:CatCh)* to generate triple transgenics. We generated the *Tg(mnx:pTagRFP)stl603* line by injecting a construct kindly shared by Dr. David McLean.

### Stochastic single cell labelling by microinjections

*Tg(eng1b:Gal4)* embryos were injected with a *UAS:Dendra* plasmid (a gift from Dr. David McLean) at the 1-2 cell stage. Final plasmid DNA concentration was 12-15 ng/µl. The embryos were transferred to system water, regularly cleaned, and allowed to develop. At 4 dpf, larvae were screened for sparse expression of Dendra in the spinal cord and selected for confocal imaging.

### Single cell labelling by electroporation

*Tg(eng1b:Gal4,UAS:GFP)* animals (4–6 dpf) were anesthetized in 0.02% MS-222 and fixed to a sylgard lined petri dish with custom made tungsten pins. One muscle segment was carefully removed to expose the underlying spinal cord. A pipette electrode (10-12 MΩ) filled with 10% Alexa Fluor 647 anionic dextran 10,000 MW (Invitrogen) in potassium-based patch internal solution, was positioned to contact the soma of the target neuron. Dye was electroporated into the cell via one to three 500 ms, 100 Hz pulse trains (1 ms pulse width) at 2–5 V (A-M systems Isolated Pulse Stimulator Model 2100). Confocal imaging was performed after 30 mins for dye filling.

### Confocal imaging

5–7 dpf larvae were anesthetized in 0.02% MS-222 and embedded in low-melting point agarose (1.5%) in a 10 mm FluoroDish (WPI). Images were acquired on an Olympus FV1200 Confocal microscope equipped with high sensitivity GaAsP detectors (filter cubes FV12-MHBY and FV12-MHYR), and a XLUMPlanFl-20x W/0.95 NA water immersion objective. A transmitted light image was obtained along with laser scanning fluorescent images to identify spinal segments. Sequential scanning was used for multi-wavelength images.

### Image analysis

Confocal images were analyzed using ImageJ (FIJI)^30^. GFP^+^ V1 neurons were marked and counted using the ImageJ Cell Counter. Segment boundaries were marked manually using the transmitted light image. For axon tracing, stitched projection images were made with the Pairwise stitching^31^ ImageJ plugin. The overlap was dictated by selecting ROIs on both images and the fused image smoothened with linear blending. Images were registered using the fluorescent channel. Number of segments traversed by V1 axons were counted manually from the stitched images.

### Electrophysiology

Whole-cell patch-clamp recordings were performed in larvae at 4–6 dpf. Larvae were immobilized with 0.1% α-bungarotoxin and fixed to a Sylgard lined petri dish with custom-sharpened tungsten pins. One muscle segment overlaying the spinal cord was removed at the mid-body level (segments 9–13). The larva was then transferred to a microscope (Nikon Eclipse E600FN) equipped with epifluorescence and immersion objectives (60X, 1.0 NA). The bath solution consisted of (in mM): 134 NaCl, 2.9 KCl, 1.2 MgCl2, 10 HEPES, 10 glucose, 2.1 CaCl2. Osmolarity was adjusted to ∼295 mOsm and pH to 7.5. To record IPSCs, APV (10 µM) and NBQX (10 µM) were also added to the bath. Patch pipettes (7–15 MΩ) were filled with either of the following two internal solutions. Current clamp (recordings of V1 spiking): (in mM): 125 K gluconate, 2 MgCl2, 4 KCl, 10 HEPES, 10 EGTA, and 4 Na2ATP. Voltage clamp (IPSCs) (in mM): 122 cesium methanesulfonate, 1 tetraethylammonium-Cl, 3 MgCl2, 1 QX-314 Cl, 10 HEPES, 10 EGTA, and 4 Na2ATP. For both solutions, pH was adjusted to 7.5 and osmolarity to 290 mOsm. Additionally, sulforhodamine 0.02% was included in the patch internal to visualize morphology of recorded cells post-hoc. Recordings were acquired using a Multiclamp 700B amplifier and Digidata 1550 (Molecular Devices). Signals were filtered at 2 kHz and digitized at 100 kHz. For IPSCs, cells were recorded at +0.3 mV (after correction for liquid junction potential of 14.7 mV.)

### Optogenetic Stimulation

A Polygon 400 Digital Micromirror Device (Mightex) was used to deliver optical stimulation. The projected optical pattern consisted of a 4×4 grid of 16 squares. Each square in the grid approximately measured 20µm x 20µm and covered 0-6 cells depending on position. One full stimulus pattern consisted of an ordered sequence of turning ON and OFF each of the 16 squares sequentially. For each small square, illumination consisted of a 20 ms light pulse (470 nm) at 50% intensity. The sequence was triggered using a TTL pulse from the Digidata to synchronize the stimulation with electrophysiology. The objective was carefully positioned over a single spinal segment prior to stimulus delivery; for each new segment, the stage was manually translated and repositioned. V1 spiking reliability was measured by delivering multiple trials to a selected square that had evoked spiking on the first trial. A similar protocol was used for all segments to obtain reliable IPSCs for measurement of conductances.

### Analysis of connectivity

Electrophysiology data were imported into Igor Pro 6.37 (Wavemetrics) using NeuroMatic^32^. Spikes and IPSCs were analyzed using custom code in Igor and MATLAB. Charge transfer for the evoked response was calculated by integrating the current in a 50 ms window from the onset of the optical stimulus (Evoked) and subtracting this from Control 1, a similar integral over a 50 ms window before the optical stimulus (Supplemental Fig. 1). This was done to account for spontaneous activity. To calculate background noise values, a similar integral for a different 50 ms window at the end of the recording (Control 2) was subtracted from Control 1 (Supplemental Fig. 1). Both the charge transfer of the evoked response and background noise were summed across the 16 squares for each segment.

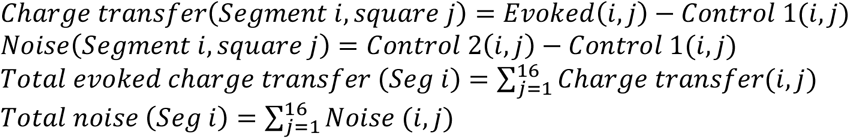

For statistical comparisons, *Total evoked charge transfer (Seg i)* was compared to *Total noise (Seg i)* for each target population using the Wilcoxon Sign Rank Test (p<0.01). The noise threshold in each Figure represents mean noise + 3*STD, averaged over all segments.

Peak amplitudes of IPSCs were calculated from the first IPSC only to avoid effects from synaptic depression/facilitation. Conductances were calculated from peak amplitude / driving force (75 mV). Input resistances was measured by an average of small hyperpolarizing pulses. Statistical tests were performed using MATLAB (R2020b, MathWorks).

### Computational modeling

Zebrafish spinal cord networks were modeled in Python as a 15-segment ipsilateral network with pacemakers located rostrally to the first segment. The Izhikevich neuron model was used to simulate individual cells in the network^33^. The following 6 parameters for each neuron were explicitly stated: *a:* recovery rate; *b:* sensitivity to spiking, *c*: reset voltage, *d*: after-spike reset rate, peak V: maximum voltage of a spike, and x: segment location. The reduced network modeled in this study included a cluster of rostrally located pacemaker neurons, V1 neurons, V2a neurons, and motor neurons (MNs). Each segment in the model incorporated a V2a neuron and a MN, with a single V1 neuron in every segment after segment 3. The network was driven using 5 electrically coupled pacemaker neurons (see Supplemental Table 1 for Izhikevich parameters) and a linear descending gradient of tonic drive to V2a neurons. Pacemakers were electrically coupled to the 6 most rostrally located V2a neurons. MNs received glutamatergic drive from V2as and glycinergic drive from V1 neurons. Electrical synapses are modeled as ideal resistors following Ohm’s law. Chemical synapses are modeled as a biexponential that accounts for rise and decay rates, as well as glycinergic and glutamatergic reversal potentials (see Supplemental Table 2 for chemical synapse parameters). V2a neurons provided descending glutamatergic projections to other V2as and V1s located up to 3 segments away. V2a neurons also made bifurcating glutamatergic projections to motor neurons, connecting to caudal motor neurons up to 3 segments away, and rostral motor neurons up to 2 segments away. V1s formed glycinergic synapses with V2as and MNs; the structure of V1 projections onto V2as and MNs was manipulated to target either local (soma within 1 to 3 segments) or distal (soma located 4 to 6 segments away). Individual weights of glycinergic synapses formed by V1s were randomized from simulation to simulation using a gaussian distribution [0.5 ± 0.25 std]; the sum of the weights of all V1 to V2a and V1 to MN was maintained between simulations. Each network configuration was simulated 15 times to generate summary data. Code for the model is available at [https://github.com/bagnall-lab/V1connecting_project].

## Results

### V1 neurons project primarily ascending axons spanning 5-10 spinal segments

V1 neurons are distributed along the length of the spinal cord^18,24^, but there are no systematic analyses of their cell numbers and morphology in zebrafish. Using confocal imaging of the *Tg(eng1b:Gal4,UAS:GFP)* fish line, we obtained cell counts of GFP+ neurons all along the length of the zebrafish larval spinal cord, which is partitioned by myotomes into ∼28 segments. V1 neurons were uniformly distributed along the rostro-caudal (R-C) axis, with an average of 18.9 ± 5.6 V1 neurons per segment (Fig. 1A, B; mean ± SD, N=10 larvae). Next, to optimize design of our subsequent mapping experiments, we investigated the extent of V1 axonal projections in the R-C axis. In mice^21^ and zebrafish^18^, V1 neurons project axons ipsilaterally and rostrally, as do their counterparts, the aINs in *Xenopus* tadpoles^22^. A subset of V1 neurons also exhibit descending axonal branches^18,34,35^. To study morphology of V1 neurons, we performed single cell labelling in the *Tg*(*eng1b:Gal4,UAS:RFP)* fish line using two approaches: single cell electroporation of fluorescently tagged dextran or micro-injection of a *UAS:Dendra* plasmid construct, followed by confocal imaging of single cells. Both techniques yielded similar results and were pooled for analysis. Fig. 1C shows an example of a representative V1 neuron labeled with *UAS:Dendra*. All V1 neurons (N=28 cells from 18 larvae) displayed an exclusively ipsilateral ascending axon, extending for a median of 6 segments. 17 / 28 neurons (60.7%) also exhibited a short descending axon branch spanning a median of 1 segment (Fig. 1D). Based on these results, we chose to build a connectivity map covering 7 segments in the ascending direction and 2 segments in the descending direction to encompass the V1 axonal extent.

**Figure 1.**
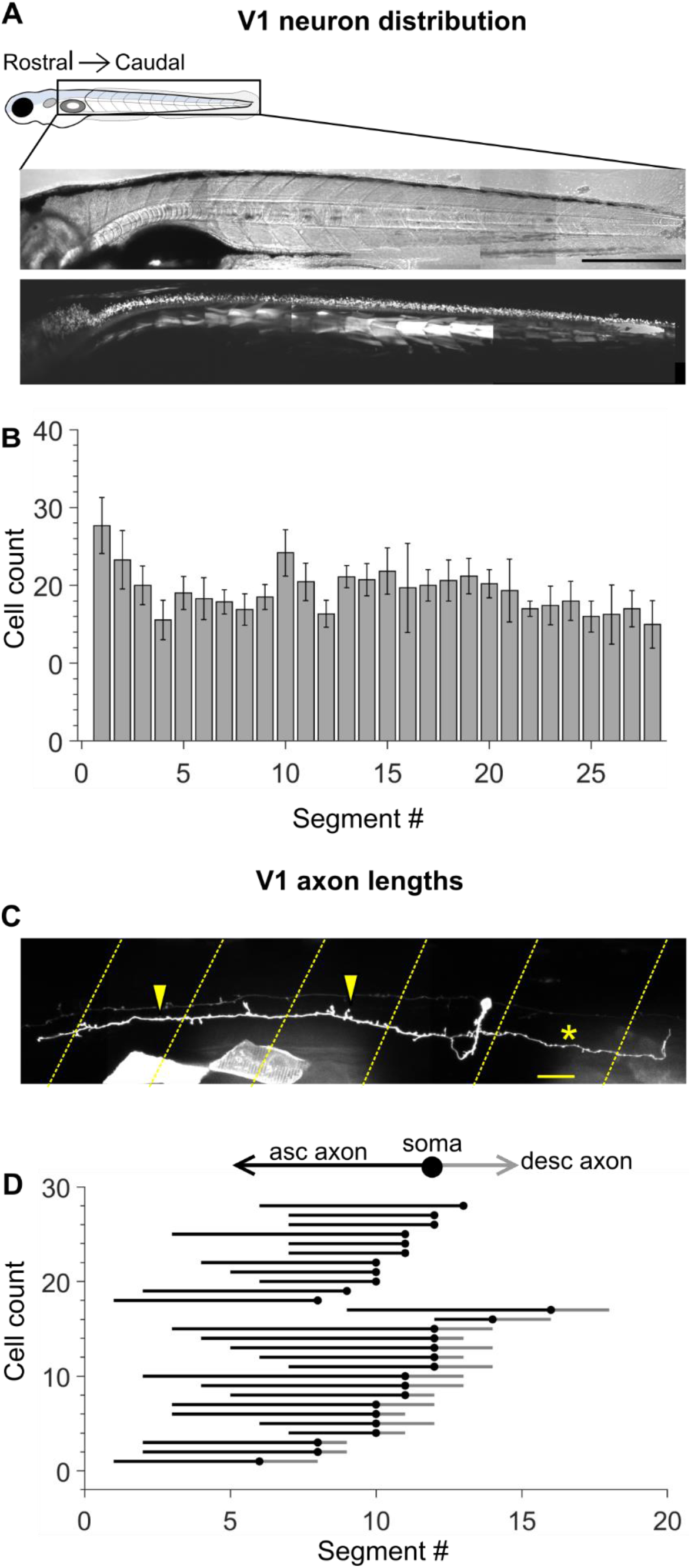
Engrailed^+^ V1 neurons project long, primarily ascending axons. A. Transmitted light image (top) and confocal image (bottom) of a 5 dpf *Tg (eng1b:Gal4,UAS:GFP)* larva. In this and subsequent Figures, rostral is to the left and dorsal to the top. Some non-specific expression of GFP is present in muscle fibers as well. Scale bar: 0.5 mm B. Bar plot showing mean cell count of V1 neurons per segment along the rostro-caudal axis. n = 15 larvae from 4 clutches. Error bars represent SEM. C. Representative example of a sparsely labelled V1 neuron in a mid-body segment. Segment borders are shown in yellow dashed lines. Arrowheads mark the ascending axon, and the asterisk marks the descending axon. Scale bar: 20 µm. D. Ball and stick plots representing the soma (ball) and ascending and descending axon lengths of V1 neurons (sticks) relative to body segments. N = 28 neurons from 18 larvae.

### Patterned optical stimulus evokes localized and reliable spiking in V1 neurons

To create a map of V1 connectivity via optical stimulation, we generated a transgenic fish line, *Tg(eng1b:Gal4,UAS CatCh)*, in which the calcium permeable channelrhodopsin CatCh^36^ was expressed in V1 neurons (schematic, Fig. 2A). We first needed to ensure that optogenetic stimulation was only effective at eliciting spiking when light was targeted near the soma of a V1 neuron, not its axon. We recorded whole cell from V1 neurons while projecting 20 x 20 µm squares of blue light via a digital micromirror device (DMD). A 4×4 grid of these squares was delivered in sequence and effectively tiled each spinal segment (Fig. 2B). The membrane potential responses of an example V1 neuron to each element of the optical stimulus is shown in Fig. 2C. Most squares elicited only subthreshold responses, but illumination of the square directly on the soma (black dot) or in a few surrounding squares effectively drove spiking (asterisks). Note that spikes recorded at the soma are very small amplitude, as previously reported for V1 neurons in zebrafish, likely reflecting their generation at some electrotonic distance from the soma^18^. Spiking elicited by illumination is represented as a heat map of spike count (Fig. 2C, right). This spatially restricted response was observed for all 10 V1 neurons recorded (Fig. 2D).

**Figure 2.**
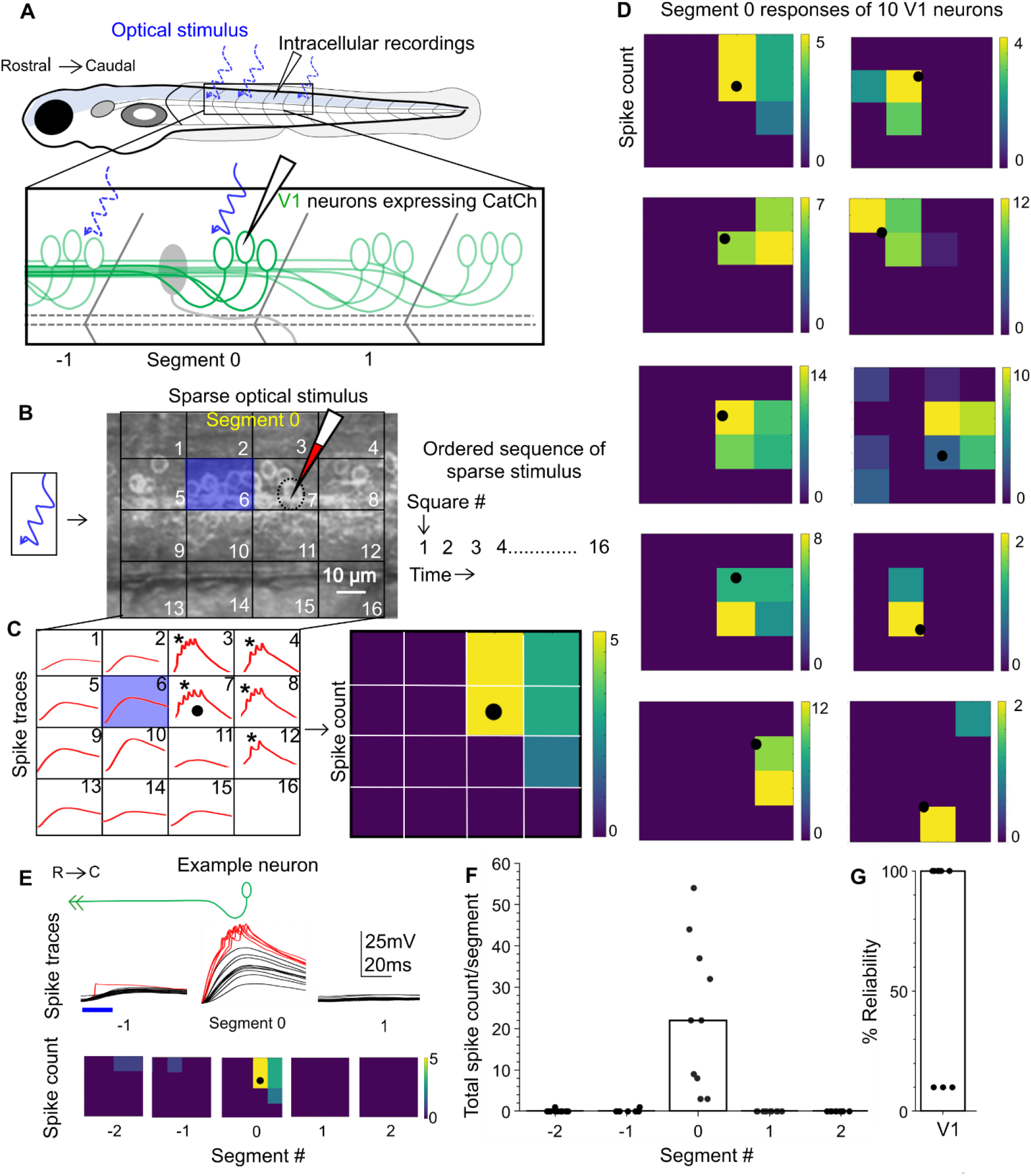
Calibration of V1 spiking with patterned optical stimulus. A. Schematic of the experimental set-up showing targeted intracellular recording and optical stimulation in *Tg(eng1b:Gal4,UAS:CatCh)* animals. B. Schematic of the patterned optical stimulus. A 4×4 grid was overlaid on approximately one segment and each square in the grid (blue square) was optically stimulated in an ordered sequence (right). Position of the recorded cell is shown as a dotted black circle. C. Illustration of the analysis. Intracellular recordings (red traces) elicited from optical stimulation in each grid square (left). Spiking is denoted by asterisks. Same data shown as a heat map and superimposed on the optical stimulus grid (right). Position of the recorded cell is indicated with a black circle. D. Heat maps generated as in C for 10 V1 neurons. E. V1 responses evoked by optical stimuli in rostral or caudal segments to the recorded neuron. Representative traces of activity (top) and spike count (bottom) of the same V1 neuron while the optical stimulation was moved along the rostro-caudal axis. Red traces indicate spiking. F. Quantification of spiking in V1 neurons as the optical stimulus is presented along the rostro-caudal axis. N = 10 neurons. Bar indicates median value. G. Reliability of spiking in these neurons with multiple trials of the same optical stimulus. Bar indicates median value.

Next, because our primary objective was to map connectivity in the R-C axis, we tested the efficacy of the optical stimulus by translating it longitudinally. An identical 4×4 illumination pattern was projected first one and then two segments away from the recorded cell, in the rostral and caudal directions. As shown in the representative example, illumination outside of Segment 0 (the recorded segment containing the V1 soma) rarely elicited any appreciable spiking responses (Fig. 2E). Antidromically evoked spiking was recorded in only 2 out of 10 cells, and even in these cases the number of spikes elicited was very low (Figs. 2E and F). Furthermore, repeated presentation of on-soma illumination reliably evoked spiking in 7 out of 10 neurons (Fig. 2G). Taken together, these data indicate that this optical stimulation is able to evoke V1 spiking only within the illuminated segment, allowing us to use this method for subsequent longitudinal mapping of connectivity.

### V1 neurons inhibit motor neurons locally

Anatomical and physiological studies indicate that V1 neurons directly inhibit motor neurons in mice^25,34,35,37^, zebrafish^3,18^, and tadpoles^22^. Based on the long ascending projections of V1 axons, we anticipated that this inhibition would extend over ∼6 segments rostrally from each V1 neuron. To examine the spatial extent of V1 inhibition, we recorded from fast primary and slow secondary motor neurons while delivering optical stimulation as above in *Tg(eng1b:Gal4, UAS:CatCh)* larvae. Primary MNs (pMNs) are identifiable by their large, laterally placed somata, low input resistances, and extensive axon arborization in characteristic patterns^38^. We targeted pMNs for whole-cell recording based on their appearance and validated their identities by post-hoc cell fills. In this and subsequent experiments, neurons were held at 0 mV in voltage clamp with a cesium-based internal solution and glutamate receptor blockers in the bath to isolate IPSCs. The patterned optical stimulus was delivered one segment at a time, caudally up to 7 segments and rostrally up to 2 segments relative to the recording site, while recording light evoked IPSCs (schematic, Fig. 3A). Fig. 3B shows representative traces of evoked IPSCs in pMNs (top) when the optical stimulus was presented 1, 3 and 7 segments caudal to the recording site, respectively. pMNs received robust IPSCs when V1s were stimulated 1 segment caudally, but surprisingly, this inhibition diminished drastically as the optical stimulation was translated further caudally (Fig. 3B, top). Charge transfer of the evoked IPSCs (Fig. 3C, inset) was calculated as the area under the curve for the 50 ms following light stimulus and compared to the noise calculated in the same way from a random post-stimulus window of the same duration (see Methods and Supplemental Fig. 1). Charge transfer for segments 0, 1, and 2 was significantly different from noise (Fig. 3B, bottom; N=26 neurons; Wilcoxon Sign Rank Test, *p* < 0.01). In contrast, the responses for segments 3, 5, and 7 were indistinguishable from noise. Charge transfer elicited by the descending axons, in Segments -1 and -2, was small in amplitude and significantly different from noise only for Segment -1. These results indicate that V1 neurons only provide appreciable inhibition onto pMNs located close to the V1 soma.

**Figure 3.**
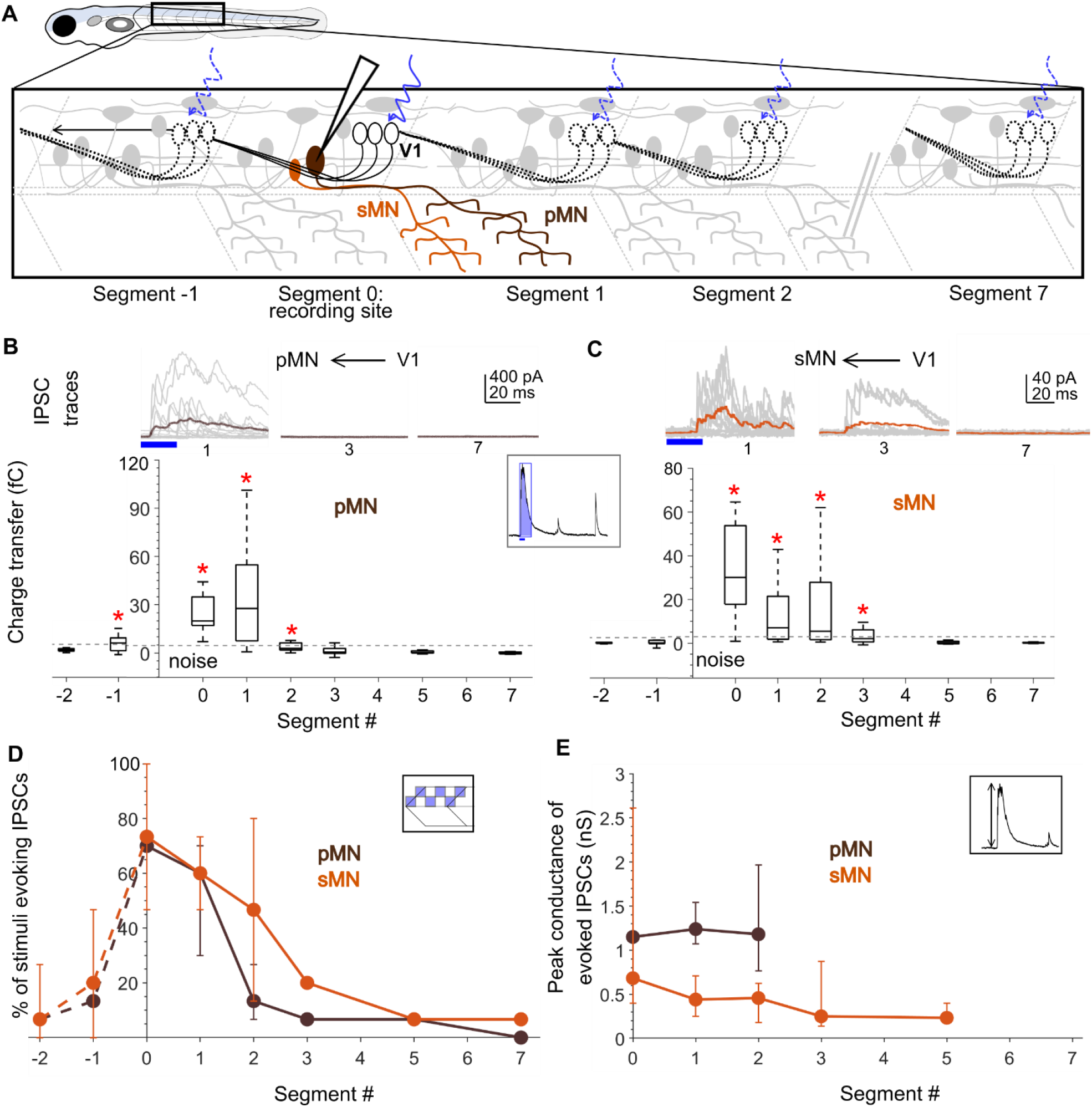
Motor neurons receive input only from local V1 neurons. A. Schematic of the experimental design showing intracellular recordings from primary (brown) and secondary (orange) motor neurons paired with optical stimulation of V1 neurons (black) along the rostro-caudal axis. B. Top: Representative overlay of 15 traces of IPSCs recorded in primary motor neurons (pMNs) during illumination of segments 1, 3, and 7 caudal to the recorded neuron position. Colored trace represents mean. Duration of the optical stimulus is shown as a blue bar. B. Bottom: Box plots showing the total charge transfer per segment (as illustrated in inset) recorded in primary motor neurons. In this and subsequent Figures, the box shows the median, 25^th^, and 75^th^ percentile values; whiskers show +/–2.7s. Dashed line indicates the level of base line noise from spontaneous activity. Red asterisks mark segments that were significantly different from noise (*p* < 0.01). N = 8-26 neurons for each data point. C. Same as in B for secondary motor neurons (sMNs). N = 10-11 neurons for each data point. D, E. Comparison of the number of squares in the optical stimuli grid that evoked IPSCs (D) and the peak conductance of IPSCs (E) in primary (brown) and secondary (orange) motor neurons. Here and in subsequent Figures, circles represent median values and error bars indicate the 25^th^ and 75^th^ percentiles. N= 8-26 pMNs and 11 sMNs.

To test V1 connectivity to slow, secondary motor neurons (sMNs), we crossed the *Tg(eng1b:Gal4,UAS:Catch)* line to a motor neuron reporter line, *Tg*(*mnx:pTagRFP*)^38^. Recordings targeted smaller motor neurons, and optical stimulation was performed the same way as above. sMNs also showed evoked IPSCs for V1 stimulation in local segments but not distally (Fig. 3C, top). Charge transfer elicited by stimulation in Segments 0-3 was significantly different from noise (N=11 neurons; Wilcoxon Sign Rank Test, *p* < 0.01), but not for distal segments 5 and 7 (Fig. 3C, bottom). Thus, similar to pMNs, sMNs are inhibited predominantly by local V1 neurons.

This decrease in evoked inhibition onto neurons located distally from the segment of stimulation could arise from a) a decrease in the number of V1 neurons connecting to motor neurons at longer distances, or b) a decrease in the strength of individual connections for distal, as opposed to proximal, synapses from V1 neurons onto motor neurons. To differentiate between these two possibilities, we first analyzed the number of grid squares that evoked IPSCs in each segment. Because V1 neurons are evenly distributed along the R-C axis (Fig. 1A, B) and our patterned stimulus uniformly covered one full segment, the number of squares evoking IPSCs can be used as a proxy for the number of connections. Fig. 3D shows a significant decrease in the percent of squares eliciting IPSCs along the R-C axis for both pMNs and sMNs (Kruskal Wallis Test and post hoc Tukey’s test, *p* < 0.01), suggesting that fewer V1 neurons in the distal segments make contact with motor neurons. As a measurement of the strength of individual synaptic connections, we analyzed the peak amplitudes of the evoked IPSCs (Fig. 3E). There was no significant difference between segments (Kruskal Wallis Test, *p* > 0.01), suggesting that the strength of individual connections is consistent along the R-C axis. Overall, these data indicate that despite projecting axons 5-10 segments rostrally, V1 neurons synapse only locally onto both primary and secondary motor neurons (< 3 segments), and that this bias in connectivity is set by the number of V1 neurons synapsing on each target, not by a change in synaptic weights.

### V2a and V2b neurons also receive inhibition locally from V1 neurons

In addition to motor neurons, premotor spinal circuits are also composed of several classes of interneurons that are crucial for setting different patterns and rhythms of movement^2^. To determine whether this pattern in V1 connectivity extends to other potential synaptic targets, we next examined their inputs onto V2a and V2b cells, which arise from a final division of the p2 progenitor class^39^. V2a (*vsx2*+, previously known as *chx10* or *alx*) neurons are glutamatergic excitatory drivers of locomotion^40–42^. V2b (GATA3+) neurons, on the other hand, are glycinergic/GABAergic and their activation slows down locomotion^28^. V1 inhibition of V2a neurons is implicated in speed control^3^, and both V2a and V2b neurons have been shown to synapse onto motor neurons^8,19,43^. We investigated the structure of V1 connectivity to these two premotor classes by crossing the *Tg(eng1b:Gal4,UAS:CatCh)* line to either *Tg(vsx2:lox-Dsred-lox:GFP)* or *Tg(gata3:lox-Dsred-lox:GFP*) to target recordings to V2a and V2b neurons, respectively^28,29^ (schematic, Fig. 4A). As shown in Fig. 4B, C, both V2a and V2b neurons could be robustly inhibited by optical stimulation of V1 neurons up to 3 segments away from the recording site. However, stimulation of V1 neurons further caudal elicited little or no inhibition. Charge transfer values for V2a neurons showed that stimulation of segments 0 - 5 evoked inhibitory responses significantly different from noise (N=14 neurons; Wilcoxon Sign Rank Test, *p* < 0.01), whereas stimulation at segment 7 did not (Fig. 4B, bottom). For V2b neurons, the effect was even more local with IPSCs elicited by stimulation at segments 0, 1, 2 and 3 significantly different from background noise (Fig. 4C, bottom; N=8 neurons; Wilcoxon Sign Rank Test, *p* < 0.01) but not at segments 5 and 7. V2a neurons also showed significant responses when V1 neurons were stimulated rostral to the recording site (Fig. 4B, Segment -2), but this was not the case for V2b neurons. As with motor neurons, we analyzed the number of squares evoking IPSCs in every segment for V2a and V2b neurons. The number of squares capable of evoking IPSCs decreased steadily as the optical stimulus was translated caudally, indicating fewer V1 neurons connecting with V2as/V2bs distally (Fig. 4E; Kruskal Wallis Test, *p* < 0.01). An analysis of the conductances of IPSCs in different segments did not show any longitudinal bias for either V2a or V2b neurons (Fig. 4F; Kruskal Wallis Test, *p* > 0.01) indicating that the differences in the connectivity along the R-C axis are shaped by a difference in the number of distally contacting V1 neurons and not by a change in the strength of the connections.

**Figure 4.**
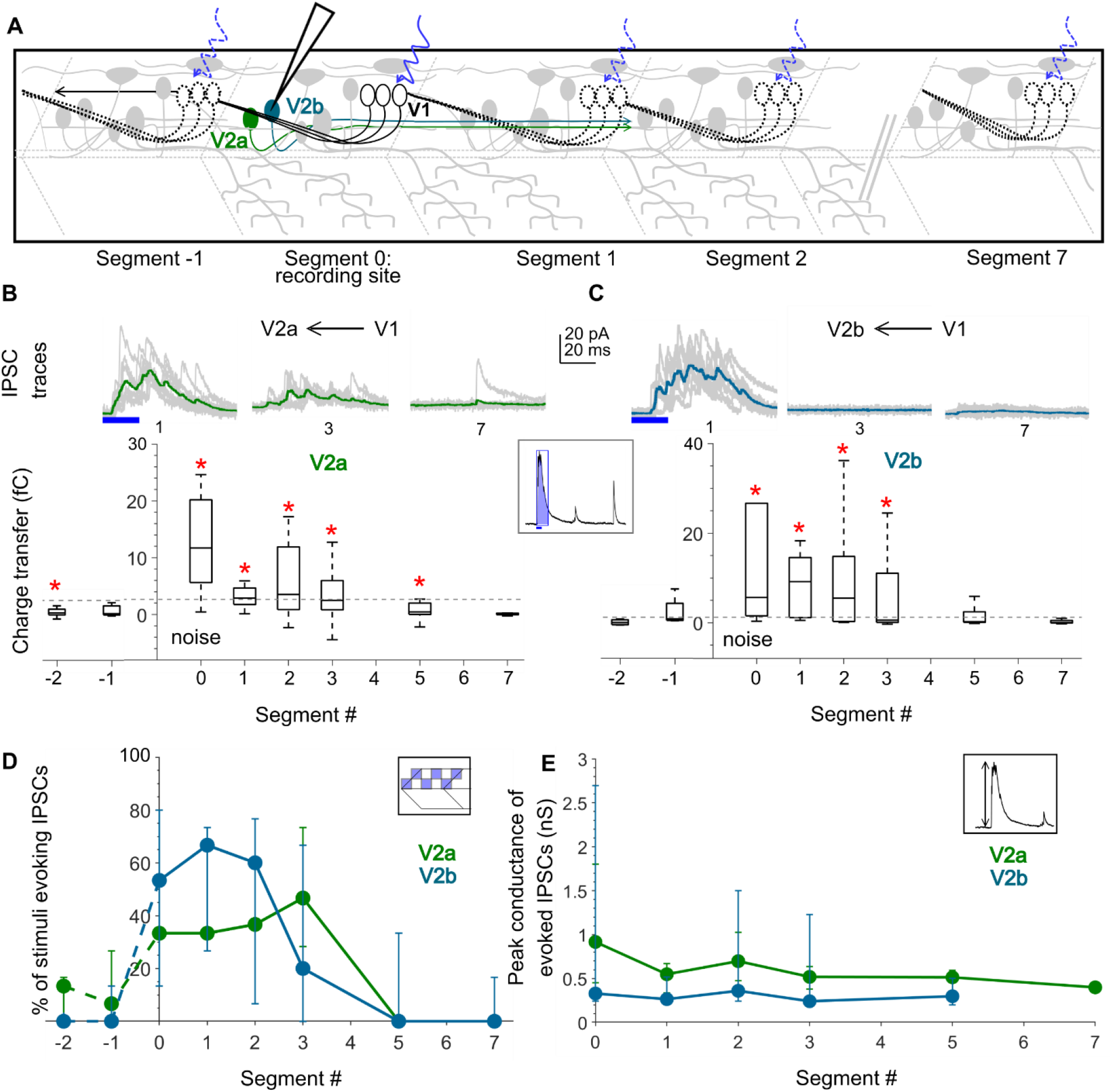
V2a and V2b neurons also receive inputs from local V1 neurons. A. Schematic of the experimental design showing intracellular recordings from V2a (green) and V2b (cyan) neurons paired with optical stimulation of V1 neurons (black) along the rostro-caudal axis. B. Top: Representative overlay of 15 traces of IPSCs recorded in V2a neurons during illumination of segments 1, 3, and 7 caudal to the recorded neuron position. Colored trace represents mean. Duration of the optical stimulus is shown as a blue bar. B. Bottom: Box plots showing the total charge transfer per segment (as illustrated in inset) recorded in V2a neurons. Dashed line indicates the level of base line noise from spontaneous activity. Red asterisks mark segments that were significantly different from noise (*p* < 0.01). N = 8-14 neurons for each data point. C. Same as in B for V2b neurons. N = 5-9 neurons. D, E. Comparison of the number of squares in the optical stimuli grid that evoked IPSCs (D) and the peak conductance of IPSCs (E) in V2a (green) and V2b (cyan) neurons. N= 8-14 V2as, 5-9 V2bs.

### CoPA neurons receive both local and distal V1 inhibition

In addition to their role in motor control, V1 neurons and their counterparts in Xenopus (aINs) are known to govern sensory gating^22^ and project to the dorsal horn^18,22,24^. In larval zebrafish, V1 neurons have been shown to directly contact Commissural Primary Ascending (CoPA) neurons^18^, a glutamatergic dorsal horn sensory population which are recruited in response to touch and cause contraversive flexion^44,45^. CoPA neurons were originally described based on their stereotypic morphology, but recent studies have identified a potential genetic marker, Mafba, a zebrafish ortholog of Mafb^46^. Based on expression of Mafb and their anatomical and functional properties, CoPA neurons are likely homologous to deep dorsal horn Laminae III/IV glutamatergic neurons arising from the dI5/dILB precursor populations that receive afferent inputs carrying innocuous mechanoreceptive signals^47,48^. During early spontaneous coiling and later in burst swimming, the CoPAs receive glycinergic inhibition that gates their activity^44^, a potential source being V1 neurons. Therefore, we next examined the longitudinal structure of V1 connectivity to CoPA neurons. CoPAs are readily distinguished by their dorsal location, large triangular somas and elongated dendrites extending several segments^45,49^, and were identified post hoc with cell fills. As above, we recorded IPSCs from CoPA neurons while delivering V1 optical stimulation along the R-C axis (Fig. 5A). Surprisingly, in contrast to our observations in motor targets, CoPA neurons received robust V1-mediated inhibition from stimulation both locally and distally, even up to 9 segments away (Fig. 5B, C, left). Charge transfer values for all segments from 0-9 were significantly different from noise (Fig. 5C, left; N=12 neurons; Wilcoxon Sign Rank Test, *p* < 0.01). No appreciable V1 connectivity was observed from the descending axonal branch of V1 neurons to the CoPAs (Fig. 5C, Segments -1 and -2).

**Figure 5.**
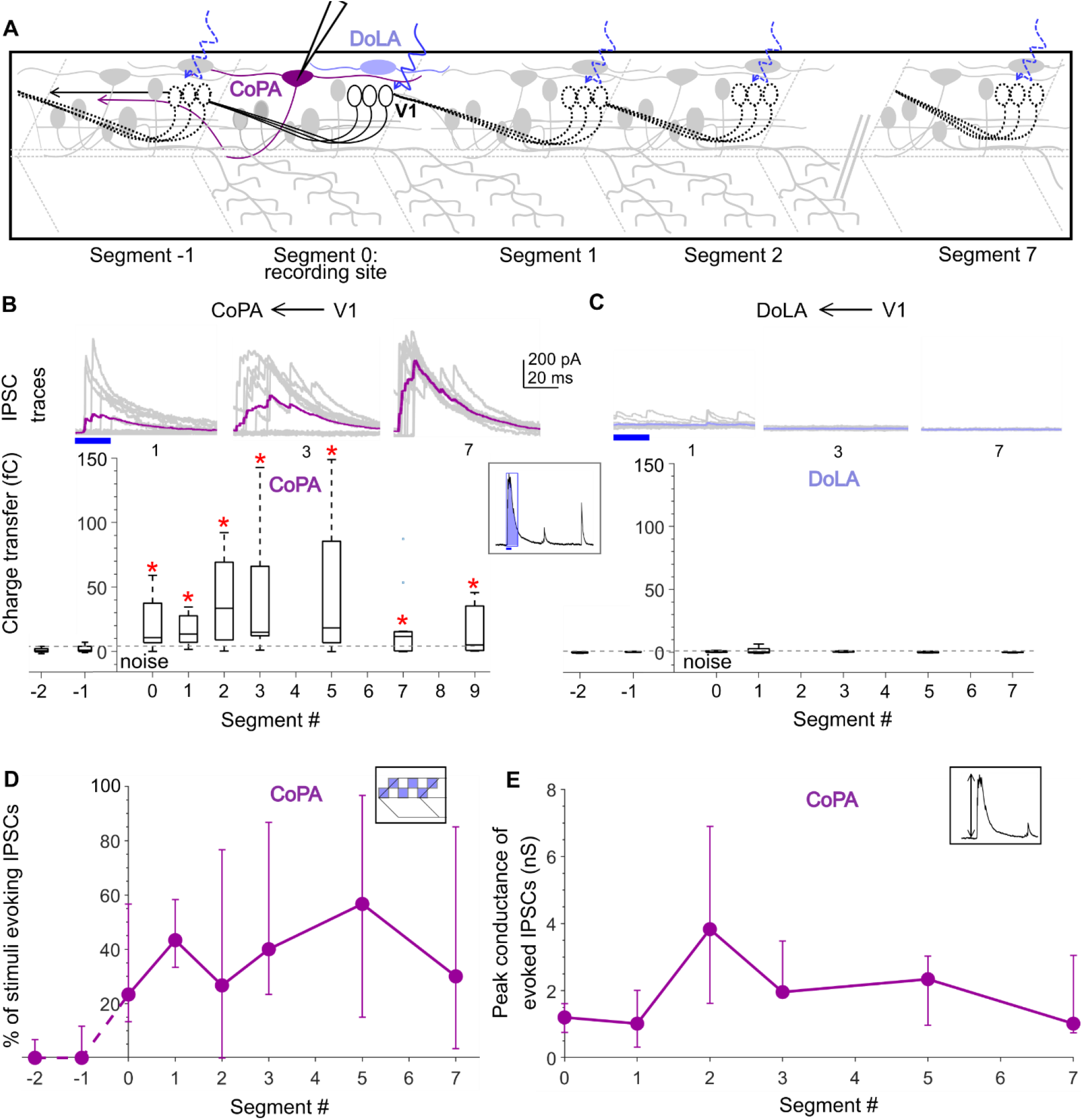
Dorsal horn CoPA neurons receive both local and long-range inputs from V1 neurons. A. Schematic of the experimental design showing intracellular recordings from CoPA (magenta) and DoLA (violet) neurons paired with optical stimulation of V1 neurons (black) along the rostro-caudal axis. B. Top: Representative overlay of 15 traces of IPSCs recorded in CoPA neurons during illumination of segments 1, 3, and 7 caudal to the recorded neuron position. Colored trace represents mean. Duration of the optical stimulus is shown as a blue bar. B. Bottom: Box plots showing the total charge transfer per segment (as illustrated in inset) recorded in CoPA neurons. Dashed line indicates the level of base line noise from spontaneous activity. Red asterisks mark segments that were significantly different from noise (*p* < 0.01). N = 7 to 12 neurons for each data point. C. Same as in B for DoLA neurons. N = 4-5 neurons for each data point. D, E. Comparison of the number of squares in the optical stimuli grid that evoked IPSCs (D) and the peak conductance of IPSCs (E) in CoPA neurons. N = 7-12 neurons.

We considered the possibility that as V1 axons ascend and travel dorsally, they connect indiscriminately to dorsal horn (sensory) targets. Therefore, we targeted another dorsal horn sensory population, the Dorsal Longitudinal Ascending (DoLA) neurons, a GABAergic population expressing Tbx16 and Islet1^50^ that is likely homologous to GABAergic Islet^+^ / dI4 cells in Lamina I-III of mouse spinal cord^51^. However, V1 neuron stimulation evoked no synaptic inputs to DoLAs, either locally or distally (Fig. 5B, C, right; N=5 cells). Therefore, the V1 connectivity to CoPAs reflects specific targeting within the dorsal horn. An analysis of the number of squares in the grid that evoke IPSCs in CoPAs revealed that there was no significant difference between local segments (Segments 0-1) and distal segments (5-7) (Fig. 5D; Kruskal Wallis Test, *p* > 0.01) indicating that a similar number of V1 neurons connect to CoPAs both locally and distally. We further compared the conductances of IPSCs between local and distal segments. No significant difference was observed between any segment (Fig. 5E; Kruskal Wallis Test, *p* > 0.01), suggesting that the strength of V1 connectivity to CoPAs is maintained all along the R-C axis.

### Other pre-motor neurons receive only local inhibition from V1 neurons

Collectively, these data indicate that V1 neurons exhibit a bias in their local vs. distal connectivity. What dictates this bias? One hypothesis is that V1 neurons connect locally to all ipsilaterally projecting targets (MNs, V2as and V2bs) but connect more broadly to contralaterally projecting targets (CoPAs). Alternatively, this bias could be based on motor related (MNs, V2as and V2bs) versus sensory (CoPAs) identities. To test these hypotheses we targeted ventral horn commissurally projecting neurons that are likely dI6/V0 identity. These neurons comprise both inhibitory and excitatory^52–54^ subsets but are characterized by a common morphological motif: a commissural, bifurcating axonal trajectory^52,53^. We utilized this distinct morphology to categorize these neurons as Commissural Pre-motor (CoPr) neurons. Neurons located dorsally to V1 neurons with commissural, bifurcating axon morphology were identified post hoc with cell fills. V1 optical stimulation was performed as before (Fig. 6A). Interestingly, CoPr neurons also received only local inhibition from V1 neurons (Fig. 6B, C). Charge transfer at Segments 0, Segment 1 and Segment 3 was significantly different from noise (Fig. 6C; N=7 neurons; Wilcoxon Sign Rank Test, *p* < 0.05) while other segments were not. We also analyzed the number of squares evoking IPSCs per segment and conductances of IPSCs. No significant differences between segments were observed for either of these parameters (Fig. 6C, D). We also examined connectivity from V1 neurons onto other V1 neurons; only at segment 0 was there significant V1-evoked inhibition (Supplemental Fig. 2). Thus, these data support the notion that V1 neurons connect locally to motor-related targets and long-range to sensory targets.

**Figure 6.**
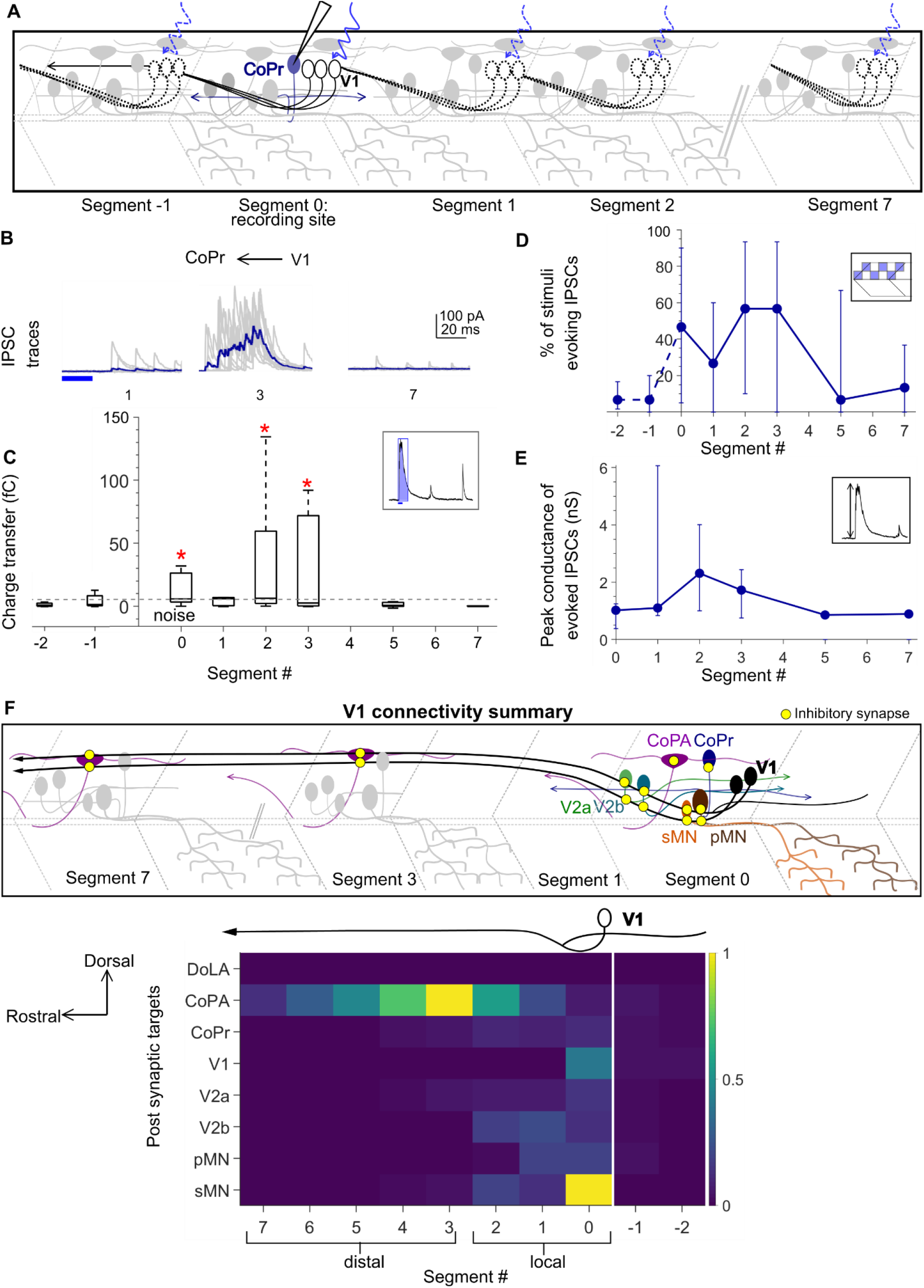
Commissural premotor (CoPr) neurons receive input from local V1 neurons. A. Schematic of the experimental design showing intracellular recording from CoPr neurons (blue) paired with optical stimulation of V1 neurons (black) along the rostro-caudal axis. B, C. Representative traces of IPSCs (B) and the total charge transfer (C) recorded in CoPr neurons with optical stimulation of V1 neurons along different segments in the rostro-caudal axis. Colored traces in B indicate mean. Dashed line in C indicates the level of base line noise from spontaneous activity. D, E. Comparison of the number of squares in the optical stimuli grid that evoked IPSCs (D) and the conductance of IPSCs (E) in CoPr neurons. N = 5 to 7 neurons for each data point. F. Summary of V1 connectivity to different post synaptic targets. Top: Schematic of the inferred connectivity of V1 neurons (black) to different targets locally and distally. Yellow dots symbolize inhibitory synapses. Bottom: Heat map showing normalized charge transfer for the different post synaptic targets along the rostro-caudal axis. The charge transfer per segment for each recorded neuronal target was normalized to its measured intrinsic neuronal conductance (inverse of Rin). Median values of normalized charge transfer for each target cell population are plotted. Values for Segment 4 and Segment 6 were interpolated as averages of the two neighboring segments. The resulting values are plotted on the same color scale for all target populations.

Fig. 6F summarizes connectivity data to all of the targets tested. The magnitude of charge transfer is not directly comparable across neurons, because inhibition’s effects will depend on its strength relative to the total conductance of the target neuron. Therefore, we normalized the charge transfer for each neuron to that cell’s intrinsic conductance (i.e., the inverse of input resistance) to compare the impact of V1 inhibition between different targets as well as across the R-C axis (Fig. 6F, bottom). We clearly observe differential connectivity from V1 neurons to sensory (CoPA) as compared to motor related (pMN, sMN, V2a, V2b and CoPr) post synaptic targets. Because sensory and motor related targets are found in the dorsal and ventral horns, respectively^2^, this heat map of V1 connectivity along the R-C axis also showed a dorsal – ventral structure. Interestingly, the contribution of the descending axonal branch of V1 neurons, though visible in the individual data sets, appeared to have minimal impact when normalized (Fig. 6F, Segments -1, -2). Taken together, these data show that although V1 neurons extend long, ascending axons spanning several spinal segments, they do not uniformly connect to all post synaptic targets along the extent of their axons. Closer to their somata (locally), V1 neurons preferentially inhibit motor and pre-motor targets. In contrast, as the axon travels rostrally, connectivity with motor and pre-motor neurons falls off sharply, and instead it inhibits sensory CoPAs (schematized in Fig. 6F, top).

### V1 connectivity to local motor populations is required for longitudinal coordination

These experimental data demonstrate that V1 connectivity is restricted to motor and premotor targets located within 1 to 3 segments. To evaluate the importance of the structure of ipsilateral inhibition on zebrafish swimming behavior, we developed a computational model of the zebrafish spinal cord. Since V1 neurons do not have any effect on left-right alternation^6,25^, we modeled only the unilateral cord. V2b neurons were excluded due to lack of knowledge on their downstream targets other than motor neurons. CoPA neurons were not included because they are thought to be active in response to unexpected touch, not during normal locomotion^44^. This reduced model comprised a cluster of pacemaker neurons and a 15 hemisegment spinal cord, consisting of MNs, V2a, and V1 neurons (Fig. 7A, see Methods for detailed description). Individual neurons were simulated with ordinary differential equations as described in the Izhikevich model^33^. V2a neurons formed glutamatergic synapses onto other V2as, V1s, and MNs, while V1s formed glycinergic synapses onto V2a and MNs. The spiking activity of MNs served as the readout of our spinal cord model.

**Figure 7.**
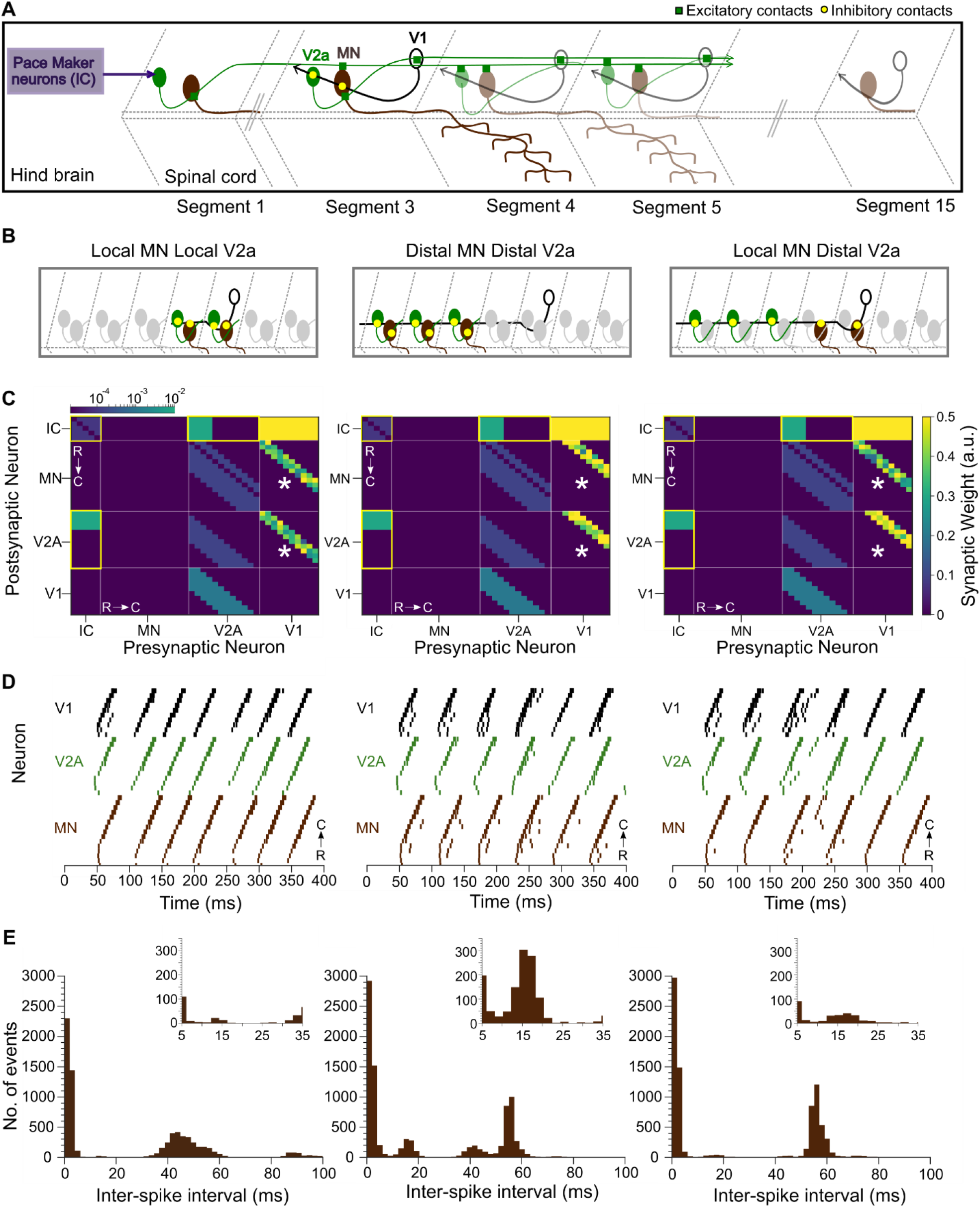
Modeling spinal circuitry with local and distal V1 circuitry. A. Schematic of the computational model showing a reduced V1 (black), V2A (green), and motor neuron (MN, brown) network driven by rostrally located pacemaker neurons (purple box). B. Schematic of 3 different network structures simulated with the network model. Left: Local MN and local V2A connectivity from V1s. V1s synapse onto V2As and MNs located within 1 to 3 segments. Middle: Distal MN and distal V2A connectivity from V1s. V1s synapse onto V2As and MNs located 4 to 6 segments away. Right: Local MNs and distal V2A connectivity from V1s. V1s synapse onto V2As located 4 to 6 segments away and MNs located within 1 to 3 segments. C. Heatmap showing connectivity weights for neurons across 3 different network models. Connections highlighted in yellow are gap junctional and follow a logarithmic scale. Asterisks indicate altered connections. D. Raster plots of spike times from 1 representative simulation of each network. E. Inter-spike interval (ISIs) frequency histograms of motor neuron spiking from 15 simulations for each network structure. Insets highlight ISIs in an intermediate range.

We first simulated a network that matched our experimental results, with local V1 inhibition (within 1 to 3 segments) onto V2as and MNs (schematic, Fig. 7B; connectivity map, 7C, left). This model recapitulated swim beats with clean rostro-caudal propagation of a locomotor wave (Fig. 7D, left). The tail beat occurred at a frequency of 21.4 Hz (mean ISI between 35 and 70 ms; 46.7 ms ± 6.1 std), as seen in an inter-spike interval (ISI) plot for all MNs (Fig. 7E, left). Next, we tested the consequences of changing V1 inhibition from local to distal by shifting V1 connections onto MNs and V2as located 4-6 segments rostrally (Fig. 7B, C, middle). The total amount of inhibition was held constant compared to the first model; only the location of the connections was altered. Distal V1 connectivity to MNs and V2as produced extraneous spikes in MNs outside of swim beats, creating erratic oscillations and aberrant “contractions” that disrupted the smooth rostro-caudal propagation (Fig. 7D, middle). To quantify these effects, we measured the ISI for each MN and found that in addition to ISIs associated with the tail beat (17.9 Hz, 55.6 ms ± 2.2 ISI, Fig. 7E, middle), new peaks in the histogram revealed MNs firing out of phase, with frequencies of 66.1 Hz (15.1 ms ± 4.4 ISI, Fig. 7E, middle) and 23.8 Hz (42 ms ± 3.2 ISI, Fig. 7E, middle). These spikes reflect “contractions” during the time immediately after passage of the locomotor wave, when normally spiking would be suppressed to avoid aberrant movement. To determine whether normal network function is impacted more by distal V1 to V2a connectivity or distal V1 to MN connectivity, we simulated a hybrid network with distal V1 to V2a but local V1 to MN connections (Fig. 7B, C, right). Interestingly, this network had fewer extraneous spikes but still exhibited slower swim beats compared to all local V1 inhibition (17.8 Hz, 56.1 ± 3.2 ms vs ∼21.4 Hz) (Fig. 7D, E, right). We also simulated the reverse hybrid model with local V1 to V2a but distal V1 to MN connections; this network exhibited a similar frequency of swim beats (21.8 Hz, 45.9 ms ± 6.1) as our experimentally derived model but produced extraneous spikes (53.3 Hz, 18.8 ms ± 7.7, Supplemental Fig. 3). Taken together, these simulations show that local V1 connectivity is crucial for normal spinal cord rhythmicity, and that alterations in this network structure affect both swim frequency and network reliability.

## Discussion

In this study, we show that V1 neurons exhibit differential connectivity to targets located proximally vs distally along the longitudinal axis of the spinal cord. Specifically, V1 neurons inhibit motor and premotor targets located nearby, and sensory targets further away, a unique connectivity pattern not described before. Furthermore, we show that this configuration has critical functional implications for propagation of the locomotor wave and R-C coordination. The results demonstrate that circuit architecture can vary along the longitudinal axis of the spinal cord, and that this architecture is important to circuit function.

### Distribution and anatomy of V1 neurons

We find that V1 neurons are evenly distributed along the R-C axis, with similar numbers per segment as reported for both V2a and V2b neurons^19,28^. In mice, not only is V1 distribution weighted to the caudal end^55^ but also transcriptionally different subsets of V1 neurons are enriched differentially in rostral versus caudal spinal segments^55,56^. It will be interesting to examine whether transcriptional profiles of V1 neurons in zebrafish reveal multiple differentially distributed subsets, or whether that is specific to the evolution of limbs. Morphologically, our observations match earlier descriptions from different amniotes. V1 neurons project primarily ascending and exclusively ipsilateral axons with a shorter descending axonal branch^18,21,22,34^. The descending axonal branch is of lower caliber and also develops later^18^. Our cell fills revealed only short descending branches (up to 2 segments), in contrast with an earlier study in zebrafish reporting longer branches up to 7 segments^14^. These morphological results were supported by physiological observations that V1-mediated IPSCs could not be elicited in recordings from targets >2 segments caudal to the stimulated segment (i.e., Segments -3 and -5; data not shown). Future work may reveal the function of long descending branches after their later development.

### Heterogeneity of V1 neurons

Recent work in mice has shown that V1 neurons can be divided into 50 transcriptionally different subtypes that exhibit distinct physiology and position in the ventral horn, implying different functions^57^. Our anatomical experiments revealed two different morphologies of V1 neurons: those with purely ascending axons (ascending V1s) and others with both an ascending axon and a descending axonal branch (bifurcating V1s) (Fig. 1C, D), suggesting the possibility of two different subclasses. Interestingly, we observed robust contacts from the descending axonal branch onto motor neurons (Fig. 3C, D) and V2as (Fig. 4C) but not with sensory CoPAs (Fig. 5C). One potential explanation is that differential connectivity to motor and sensory targets is accomplished by two different V1 subclasses; i.e., ascending V1s project long distances and connect only to sensory CoPAs whereas bifurcating V1s only project locally and connect to motor targets. However, our data do not support this hypothesis: there are no differences in the ascending axon trajectories (axonal length and D-V positions of the axons) or even the D-V position of the somas between these two subtypes. Therefore, we conclude that individual V1 neurons likely connect to both sensory and motor targets differentially along their projections.

In limbed vertebrates, multiple functional subclasses of V1 neurons have been identified: Renshaw cells involved in feedback control, and Ia inhibitory neurons participating in flexor-extensor reciprocal inhibition^58^. Both these subclasses contact motor neurons but receive different inputs^59^. Since our data maps the output of V1 neurons, and not the input, we cannot evaluate whether V1 neurons in our study are similar to Renshaw cells or Ia inhibitory neurons. Future delineation of subtypes of V1 neurons in zebrafish, including analysis of their inputs from motor neuron collaterals^60^, will help elucidate additional conserved functions of these neurons in motor control.

### V1 influence on speed regulation

Ablating or inhibiting V1 neurons results in slower speeds of locomotion in both mouse and zebrafish^3,6,23^. V1 neurons appear to govern speed through two different mechanisms: suppression of spiking in slow motor neurons and burst termination in fast motor neurons. In accordance with this, we also observed robust evoked IPSCs from V1 neurons to fast, pMNs (Fig. 3C). Although the magnitude of evoked IPSCs was higher in fast pMNs (Fig. 3C), the extent and impact of V1 inhibition was greater in slow sMNs after normalization to conductance (Fig. 6F), consistent with the idea that V1 neurons suppress slow MNs to permit fast swim^3^. Our results also confirm that V1 neurons directly inhibit V2a neurons^3^. Moreover, results from our model indicate that local connectivity to both motor neurons as well as V2as is necessary to maintain fast speeds of locomotion and R-C propagation of the locomotor wave (Fig. 7). Another spinal population affecting locomotor speed is the V2b class, an inhibitory, ipsilaterally projecting interneuron population. Loss of V2b neurons result in an increase in locomotor speed, suggesting that V2bs act as brakes on locomotion^28^. Direct inhibition of V2b neurons by V1 neurons, resulting in disinhibition of motor neurons, could be yet another mechanism by which V1 neurons facilitate high locomotor speeds. Future analysis with models including different speed modules as well as V2b inhibition will help shed light on the fine control of these various mechanisms of speed regulation.

Like motor neurons, V1 neurons themselves can be categorized into fast and slow subtypes based on the speed at which they get recruited. Dorsal V1 neurons are recruited at slow locomotor speeds compared to more ventral V1 neurons that are recruited at faster speeds, opposite to the D-V organization of motor and excitatory neurons^61^. We analyzed V1-evoked charge transfer based on the D-V position of the V1 optogenetic stimulus but did not find any clear relationship between the D-V position of V1 stimulation and connectivity with fast or slow motor neurons (data not shown). However, the optogenetic stimulus activated V1 neurons in adjacent positions (Fig. 2D), and therefore this result is not conclusive.

### Effects on rostro-caudal coordination

A model of longitudinal coordination in Xenopus demonstrated a requirement for rostrally biased distribution of excitatory neurons as well as ascending excitation for normal locomotor propagation^62^. This model did not feature any ipsilateral inhibition, though separate studies have shown that R-C coordination required both excitatory and inhibitory spinal pathways to be intact^15^. Our data for the first time point to a major role of V1-mediated ipsilateral inhibition in R-C coordination. Even though the total amount of inhibition was kept similar, only short range V1 inhibition to both excitatory V2a neurons and motor neurons was able to produce reliable propagation of the locomotor wave. Taken together these studies suggest that there is more than one mechanism at play for executing R-C propagation. In our model, we did not factor in ascending V2a subsets, rostral biases in neuron distribution or contralateral influences. In future, it will be interesting to build a complete model of the spinal cord to see how these different mechanisms interact.

### Impact of V1 connectivity to sensory functions

Our results show robust inhibition of sensory CoPA neurons by V1 neurons, in agreement with previous observations in zebrafish^18^ and Xenopus counterparts^22^. V1 neurons in mouse also project to the deep dorsal horn^24^, but their specific targets and functions have not been elucidated. Additionally, we show that V1 neurons connect with CoPAs all along their axonal arbors (up to 7-9 segments). Long range suppression of CoPAs indicates that broad impact is the goal of V1-mediated inhibition of sensory responses. In contrast, V1 inhibition of motor targets is local, reflecting precision in timing relative to other segments. Inhibition onto CoPA neurons is thought to be shunting, altering the neuron’s membrane resistance and reaction to subsequent excitatory inputs^44^. This is different from the hyperpolarizing inhibition seen in the case of motor neurons^3^. These results would imply that not only is the spatial pattern of V1 inhibition different between sensory CoPAs and motor neurons but also the physiological effect. It will be interesting in future to see what cellular compartments are targeted by V1 neurons in these two sensory and motor targets to further understand how the structure of connectivity can impact function.

### Other examples of differential connectivity

In this study we show that V1 axons target different post-synaptic populations as they traverse the length of the spinal cord. Other types of projection neurons are reported to contact different classes of neurons along their axonal projections, but only because they travel to multiple distinct regions of the nervous system (for example, corticospinal neurons projecting to both pons and spinal cord^63^). In contrast, cortical pyramidal neurons target similar postsynaptic partners regardless of whether they are synapsing locally or long-range^64^. The differential local to long-range connectivity within one region that we describe here appears unusual; however, it might be a common strategy in the spinal cord due to the propagation of locomotor activity that requires temporal patterning of spinal activity in the R-C axis. In the spinal cord, modular organization of motor neurons and interneurons develops sequentially^61,65^. It will be interesting in future to study how the differential targeting of V1 neurons is set up developmentally and how this corresponds to the emergence of different behaviors like the touch reflex and locomotion.

The diversity of spinal interneurons, their organization, and myriad functions continue to pose challenges in understanding spinal circuits. Even for a relatively simple organism like larval zebrafish, we describe a complex pattern of connectivity within the same neuronal class, suggesting a primitive and intricate code buried within these spinal circuits. Our results further demonstrate significant interconnectivity between spinal interneuron populations, an area that requires future characterization to help decipher the underlying neuronal code. Future studies aimed at exploring other dorsal horn targets and analysis of V1 subsets in zebrafish will help understanding the role of V1 neurons in encoding sensory-motor control.

## Acknowledgments

We would like to thank Dr. David McLean and Dr. Sandeep Kishore for kindly providing us with the *mnx:pTagRFP* construct and Dr. Rich Roberts for helping create the *Tg(mnx:pTagRFP)stl603* fish line. This research was funded by the Pew Scholar Award (M.W.B), R01 DC016413 (M.W.B), a McKnight Scholar Award (M.W.B.), and the McDonnell Center for Cellular and Molecular Neurobiology Postdoctoral Fellowship 2021 (M.S.).

---

**Supplemental Figure 1:**
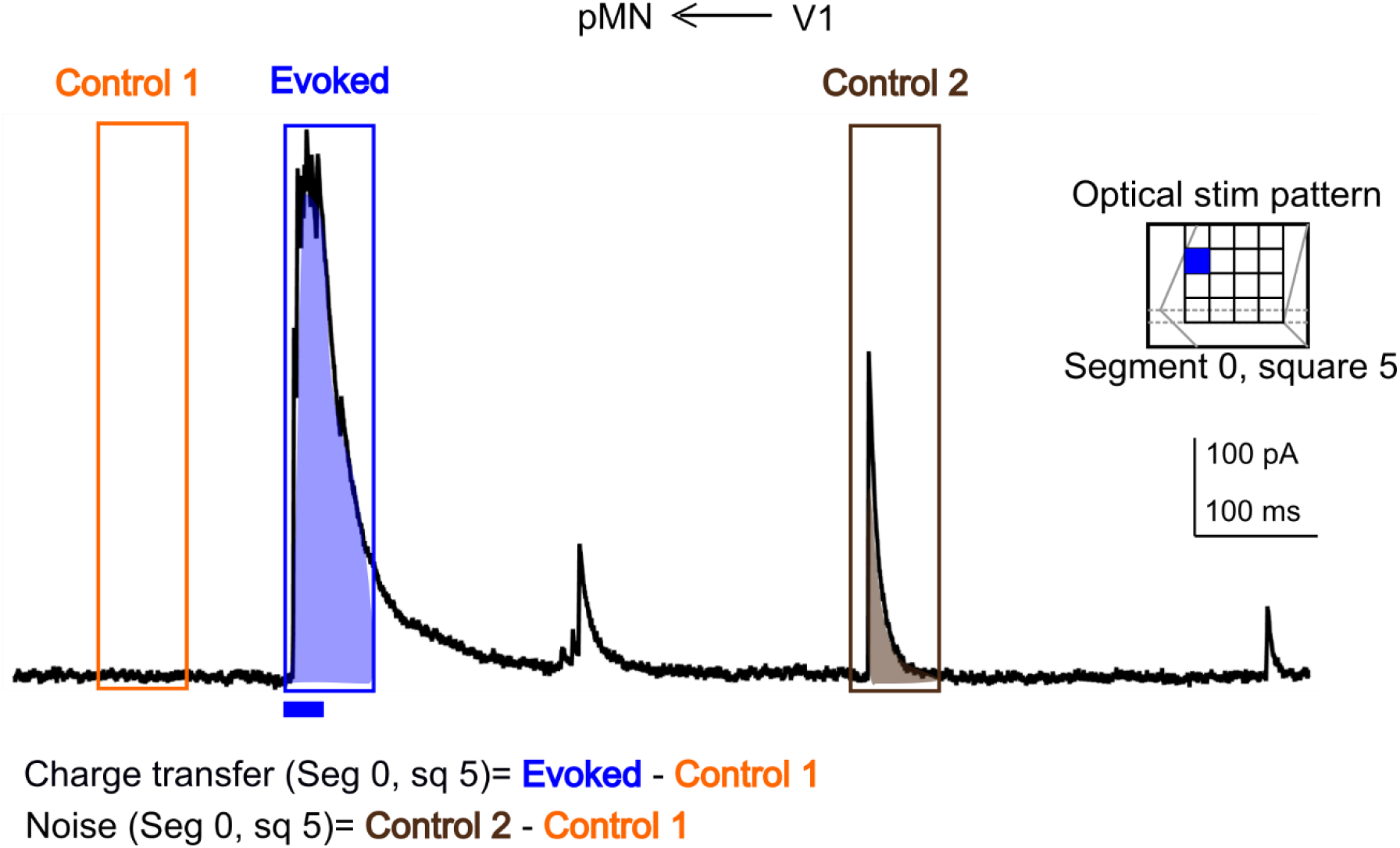
Analysis of charge transfer per segment and calculation of noise. Representative trace of IPSCs recorded in a single primary motor neuron during illumination of square 5 in Segments 0 (inset). Duration of the optical stimulus is shown as a blue bar. Charge transfer for the evoked response was calculated by integrating the current in a 50 ms window from the onset of the optical stimulus (Evoked, shaded region in the blue block). To account for spontaneous activity, the Evoked response was subtracted from Control 1, a 50 ms window before the onset of the stimulus (Orange block). To calculate noise, the charge transfer in a separate 50 ms window post stimulus was calculated (Control 2, shaded region in the brown block) and this was subtracted from Control 1. Charge transfer (Seg 0, sq 5) = Evoked - Control 1 Noise (Seg 0, sq 5) = Control 2 - Control 1 Total evoked charge transfer (Seg 0) = Charge transfer (Seg 0, sq 1) + Charge transfer (Seg 0, sq 2) ………+ Charge transfer (Seg 0, sq 16) Total noise (Seg 0) = Noise (Seg 0, sq 1) + Noise (Seg 0, sq 2) + Noise (Seg 0, sq 16) For statistical comparisons, total charge transfer per segment was compared to total noise per segment for each neuron.

**Supplemental Figure 2:**
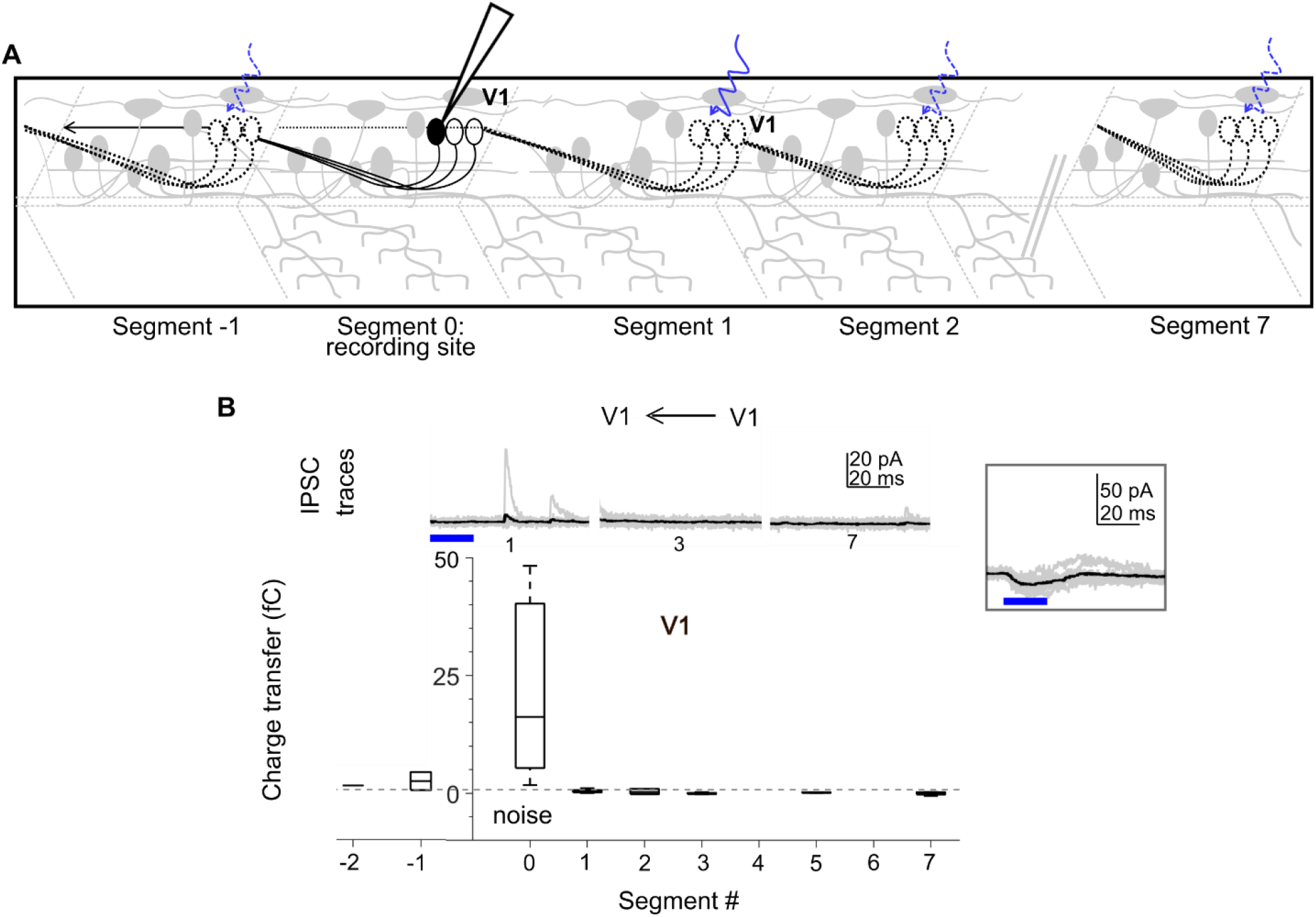
V1 neurons inhibit other V1 neurons only locally. A. Schematic of the experimental design showing intracellular recordings from V1 neurons paired with optical stimulation of V1 neurons along the rostro-caudal axis. B. Top: Representative overlay of 15 traces of IPSCs recorded in V1 neurons during illumination of segments 1, 3, and 7 caudal to the recorded neuron position. Colored trace represents mean. Duration of the optical stimulus is shown as a blue bar. B. Bottom: Box plots showing the total charge transfer per segment recorded in V1 neurons. Dashed line indicates the level of base line noise from spontaneous activity. N = 5 neurons for each data point. Inset: Recording from the same cell holding at -65 mV showing slow CatCh induced depolarization. Blue bar represents duration of the optical stimulus.

**Supplemental Figure 3:**
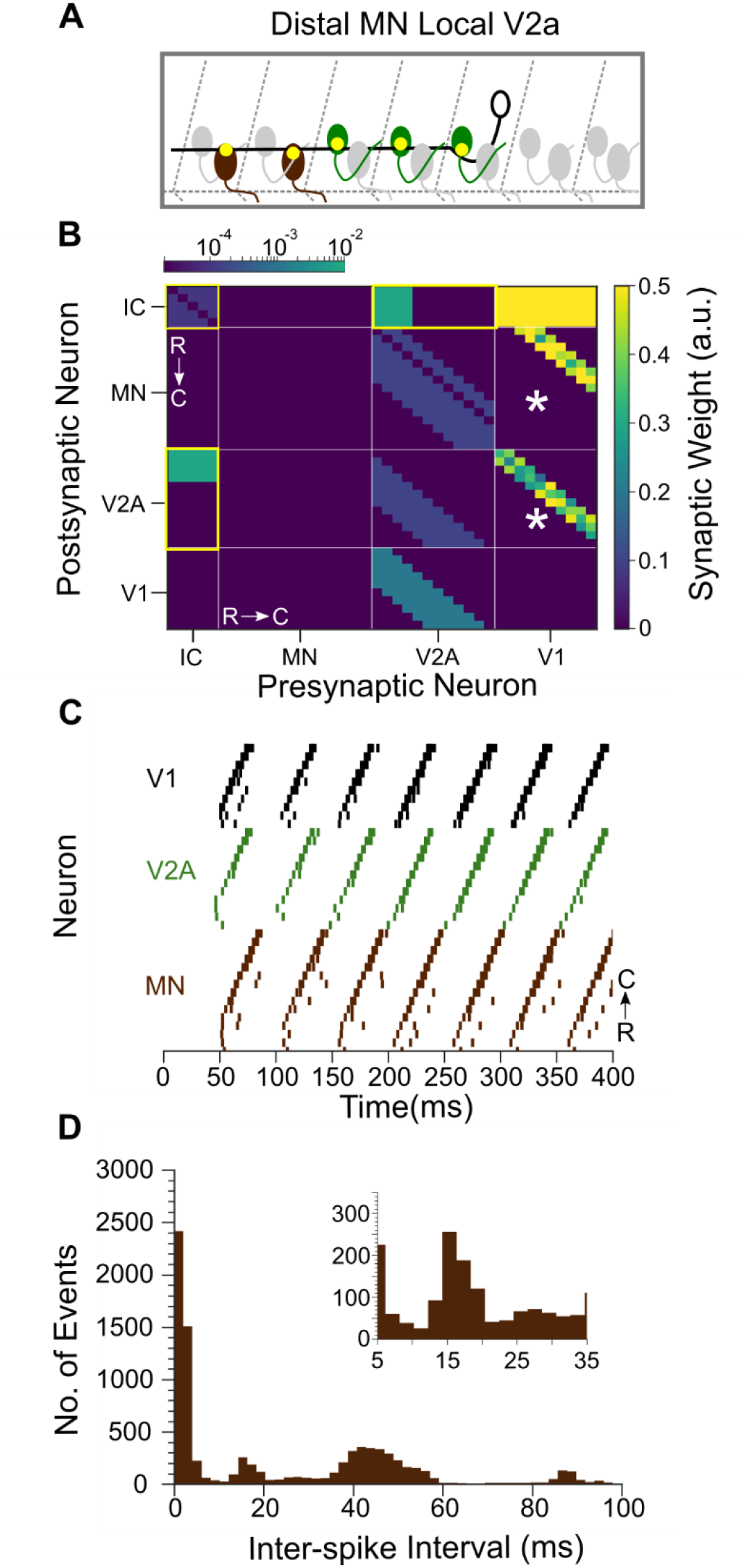
Spinal cord model with local V1 to V2a and distal V1 to MN connectivity. A. Schematic showing local V2a and distal MN connectivity from V1s. V1s synapse onto MNs 4 to 6 segments away and V2as located within 1 to 3 segments. B. Heatmap showing connectivity weights (scale bar, right). Connections highlighted in yellow are gap junctional and follow a logarithmic scale (top). Asterisks highlight the portion of the model that was altered, with connectivity shifted from more local to distal (more rostral) positions. C. Raster plots of spike times from 1 representative simulation. Rostrally located neurons are at the bottom within each neuron class, and thus the locomotor propagation moves from bottom to top. E. Inter-spike Interval (ISIs) frequency histograms of motor neuron spiking from 15 simulations of this network structure. Inset highlights ISIs from 5-35 ms.

**Supplemental Table 1:** Izhikevich Parameters

| Table 1: Izhikevich Parameters |  |  |  |  |  |
| --- | --- | --- | --- | --- | --- |
| Neuron Type | a | b | c | d | Peak V (mV) |
| Pacemakers | 0.02 | 0.25 | -50 | 2.0 | -10 |
| V2a | 0.1 | 0.25 | -53 | 6.0 | 0 |
| V1 | 0.2 | 0.25 | -53 | 6.0 | 0 |
| MN | 0.1 | 0.25 | -53 | 6.0 | 0 |

**Supplemental Table 2:** Chemical Synapse Parameters

| Table 2: Chemical Synapse Parameters |  |  |  |
| --- | --- | --- | --- |
| Synapse Type | $\tau$ rise | $\tau$ decay | Reversal V (mV) |
| Glycinergic | 0.5 | 3.0 | -70 |
| Glutamatergic | 0.25 | 1.0 | 0 |

## Notes

### Competing Interest Statement

The authors have declared no competing interest.

